# In Vitro Potentiation of Carbapenems with Tannic Acid Against Carbapenemase Producing Enterobacteriaceae: Exploring Natural Products as Potential Carbapenemase Inhibitors

**DOI:** 10.1101/366104

**Authors:** Anou M. Somboro, John Osei Sekyere, Daniel G. Amoako, Hezekiel M. Kumalo, René Khan, Linda A. Bester, Sabiha Y. Essack

## Abstract

Resistance to antibiotics is increasing worldwide, necessitating urgent action to sustain the efficacy of existing antibiotics in clinical use. We show that tannic acid (TA) in combination with carbapenems can reduce and/or reverse the minimum inhibitory concentrations (MICs) of carbapenems to susceptible values in Enterobacteriaceae that express class A and B carbapenemases. MICs of carbapenems in the presence and absence of TA and other efflux pump inhibitors, TA-carbapenemases inhibition assays and computational studies were undertaken to determine the effect of TA on carbapenem susceptibility in *Enterobacteriaceae*. TA had the greatest effect on metallo-β-lactamases (MBLs) followed by class A serine-β-lactamases (SBLs). Antibiotic susceptibility testing showed that TA reversed the MICs of MBLs to susceptible values whilst substantially reducing the MICs of SBLs (class A). Tolerable cytotoxicity effect was observed for the concentrations tested. TA inhibited enzymes with a marked difference between ≈50% inhibition (IC_50_) for NDM-1 and KPC-2. Computational studies including molecular docking, molecular dynamics simulations and binding free energy calculations showed that TA interact with both MBLs and SBLs hydrophobic sites. Moreover, TA had a stronger binding affinity for MBLs than SBLs as the MBLs, specifically VIM-1 and NDM-1, interact with a larger number of their catalytic active-site residues than that of OXA- 48 and KPC-2. These *in vitro* evaluations together with computational simulation explain the potentiating effect of TA toward carbapenems against carbapenem-resistance enterobacteriaceae. This study proposes TA as a promising adjuvant for MBLs and SBLs.

## 1. Introduction

Antibiotic-resistant infections are increasing morbidities and mortalities worldwide (Powledge 2004). The exponential rise in the prevalence and global dissemination of carbapenem-resistant and carbapenemase-producing *Enterobacteriaceae* (CRE and CPE, respectively) in communities, hospitals, and the environment is of grave concern as carbapenems are considered “last-resort antibiotics” for difficult-to-treat multi-drug resistant infections (Sekyere et al. 2016; Osei Sekyere, Govinden, Bester, et al. 2016; Yang et al. 2016). The World Health Organization (WHO) recently listed CRE/CPE as “critical pathogens” which are currently the biggest threat to healthcare due to their multidrug resistance to all known classes of antibiotics. Therefore, there is the need to develop novel and efficient strategies to overcome the WHO global priority pathogen list (PPL) (World Health Organization 2017).

Several intervention strategies have thus far been proposed to overcome antibiotic resistance, including the inhibition of mutations, sequential treatment, and use of antisense oligomers to prevent expression of the *acrA* gene that forms part of the *acrAB* efflux pump, which mediates multidrug resistance (Ayhan et al. 2016). As novel antibiotics are bound to also become ineffective over time, it is imperative to sustain the efficacy of existing antibiotics. In this context, the use of adjuvants that inhibit efflux and hydrolysing enzymes of antibiotics are gaining widespread interest (Ayhan et al. 2016; Sekyere & Amoako 2017). For instance, Chusri et al. (Chusri et al. 2009) and Tintino et al. (Tintino et al. 2016) showed that tannic acid (TA) a type of polyphenolic biomolecule which acts as an efflux-pump inhibitor (EPI) and potentiates the effects of several non-β-lactam antibiotics in *Acinetobacter baumannii* and *Staphylococcus aureus*, respectively. Payne et al. also characterized series of tricyclic natural product-derived as potent large-spectrum MBL inhibitors (Payne et al. 2002). Whilst plant-based agents or extracts are perceived as safe, none are currently approved for clinical use. This study herein characterized TA, a plant-derived polyphenol compound for its property to protect carbapenems from MBLs and SBLs hydrolysis.

Using a large international collection of clinical CPEs of diverse species that express known class A, B, and D carbapenemases (Nordmann et al. 2012; Osei Sekyere & Amoako 2017; Sekyere & Amoako 2017), we show herein for the first time, to our knowledge, that TA reduces class A and B carbapenemase-mediated carbapenem resistance. We show that this reduction in MICs is due to inhibition of carbapenemase activity in the CPE isolates, which was supported by enzymatic assay and computational simulation. This study also showed that TA does not act as an efflux pump inhibitor (EPI) in Enterobacteriaceae contrary to what was reported in *A. baumannii* and S. *aureus* (Chusri et al. 2009; Tintino et al. 2016).

## 2. Materials and Methods

### 2.1. Bacterial strains, efflux pump inhibitors (EPIs), and tannic acid (TA)

Seventy-two CRE isolates comprising 46 clinical isolates obtained from private hospitals in South Africa (Osei Sekyere, Govinden, & Essack 2016; Osei Sekyere & Amoako 2017; Sekyere & Amoako 2017) and 26 reference isolates of international origin obtained from Institut National de la Santé et de la Recherche Médicale (U914), Paris, France, (Nordmann et al. 2012) were used in this study. The species breakdown of the 72 *Enterobacteriaceae* isolates are shown in Tables 1-3; Tables S1-S2 and S4. *E. coli* ATCC 25922 was used as control in all experiments. Certified pure products of meropenem (MEM), imipenem (IPM), tannic acid (TA), carbonyl cyanide m-hydrophenylhydrazine (CCCP), reserpine (RSP), verapamil (VRP), thioridazine (TZ), and chlorpromazine (CPZ) as well as cation-adjusted Mueller-Hinton broth and agar were purchased from Sigma Aldrich (St. Louis, MO, USA). VRP, RSP, TZ, and CCCP’s ability to inhibit efflux activity have been already described (Sekyere & Amoako 2017).

**Table 1:**
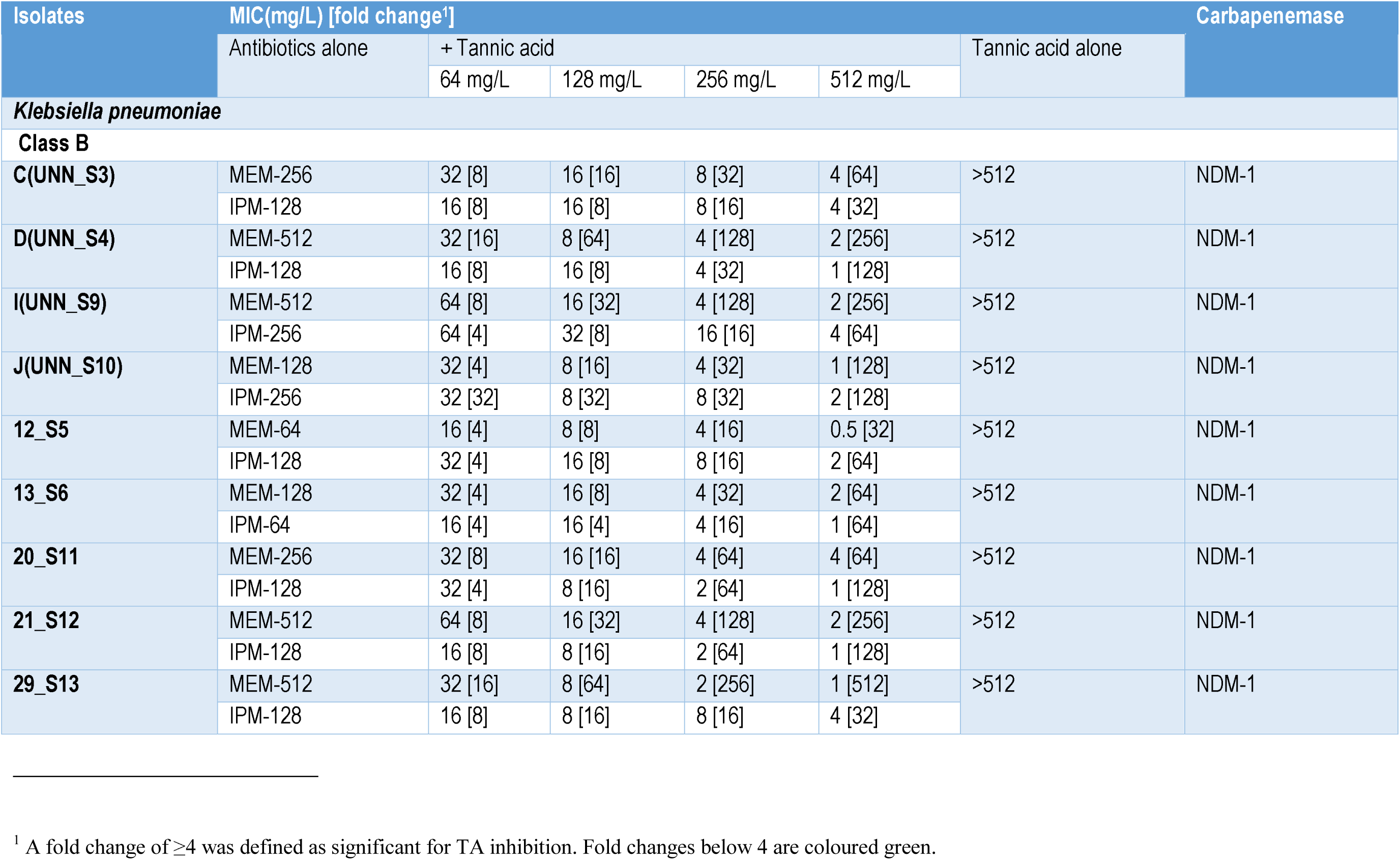
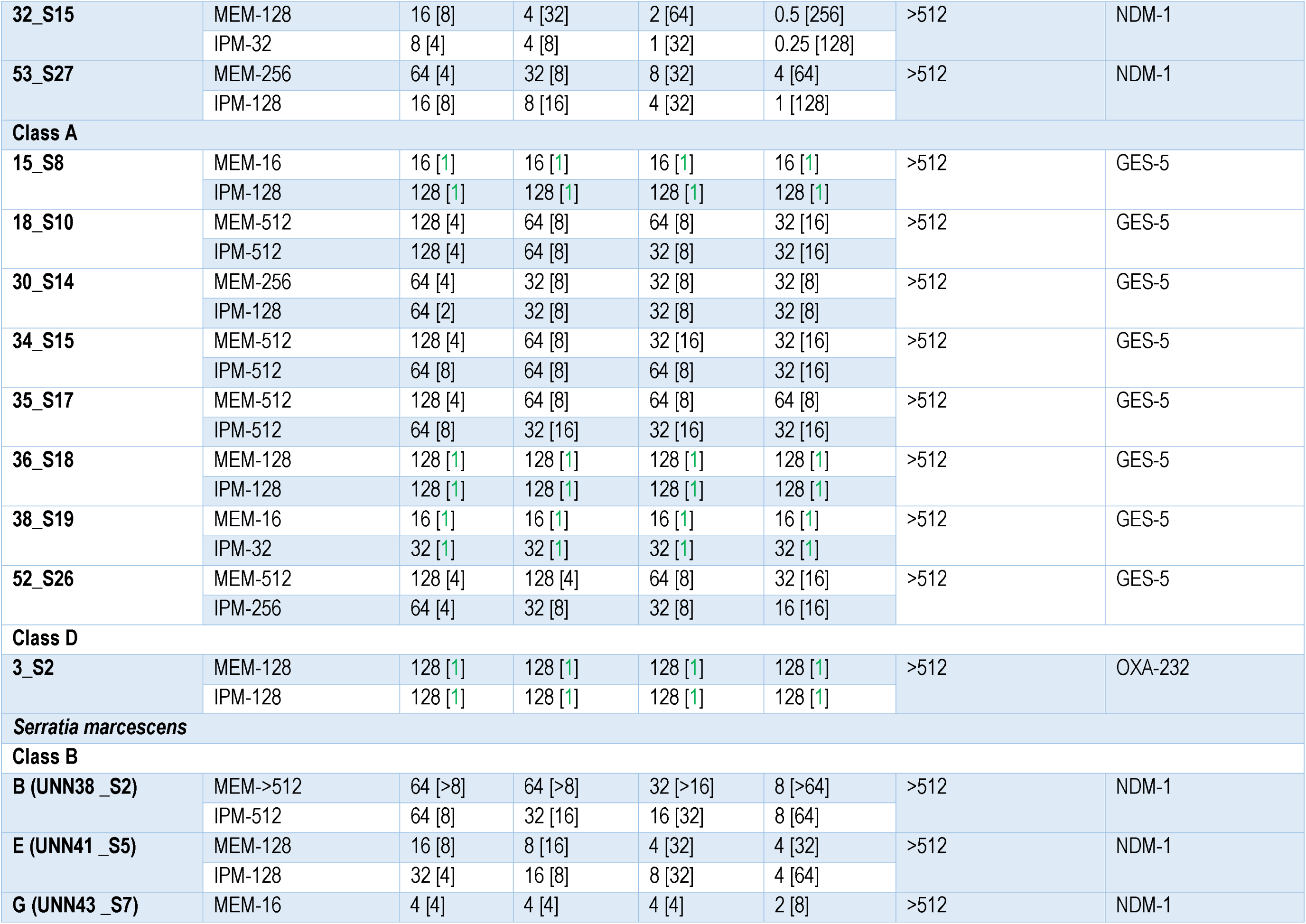
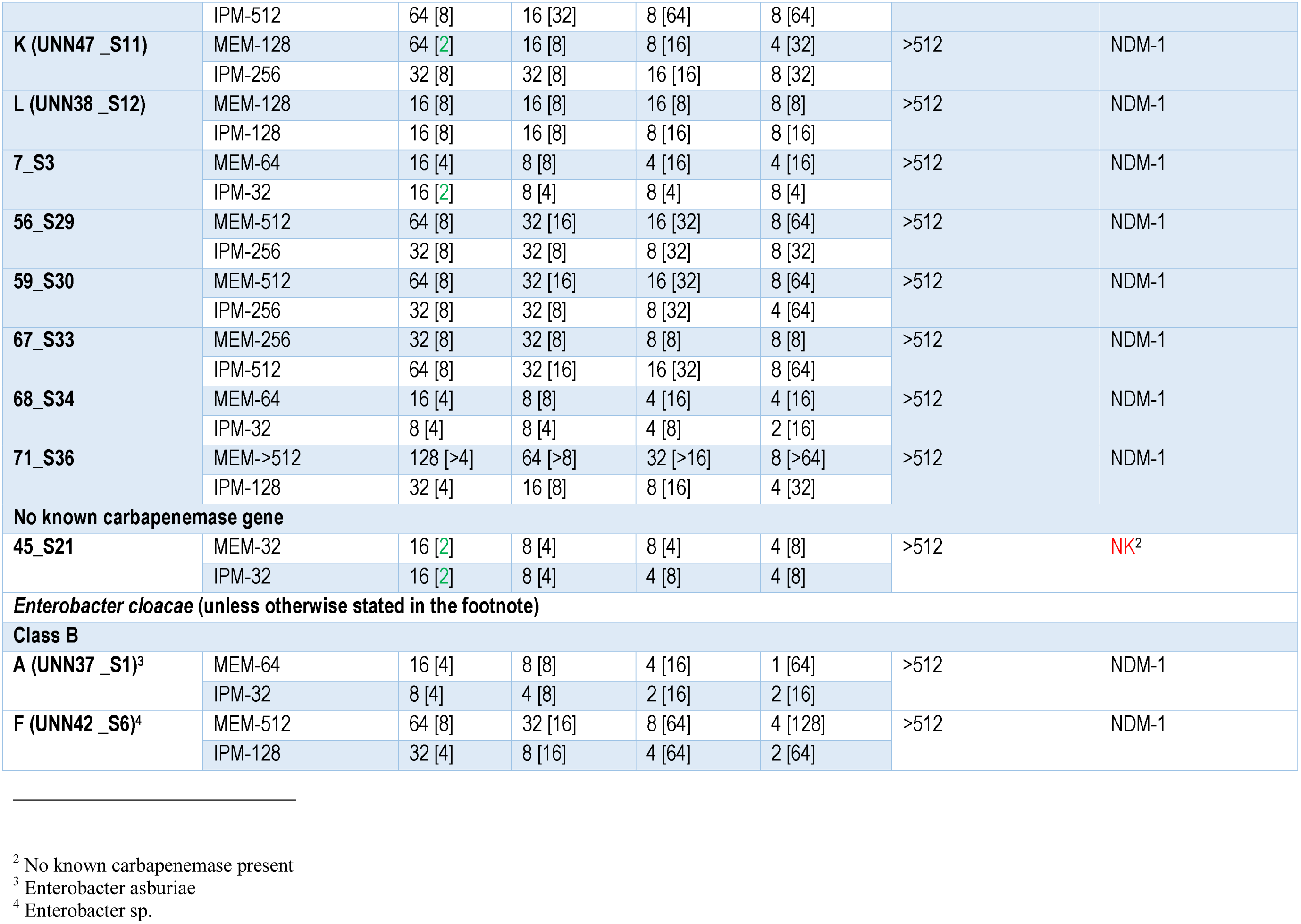
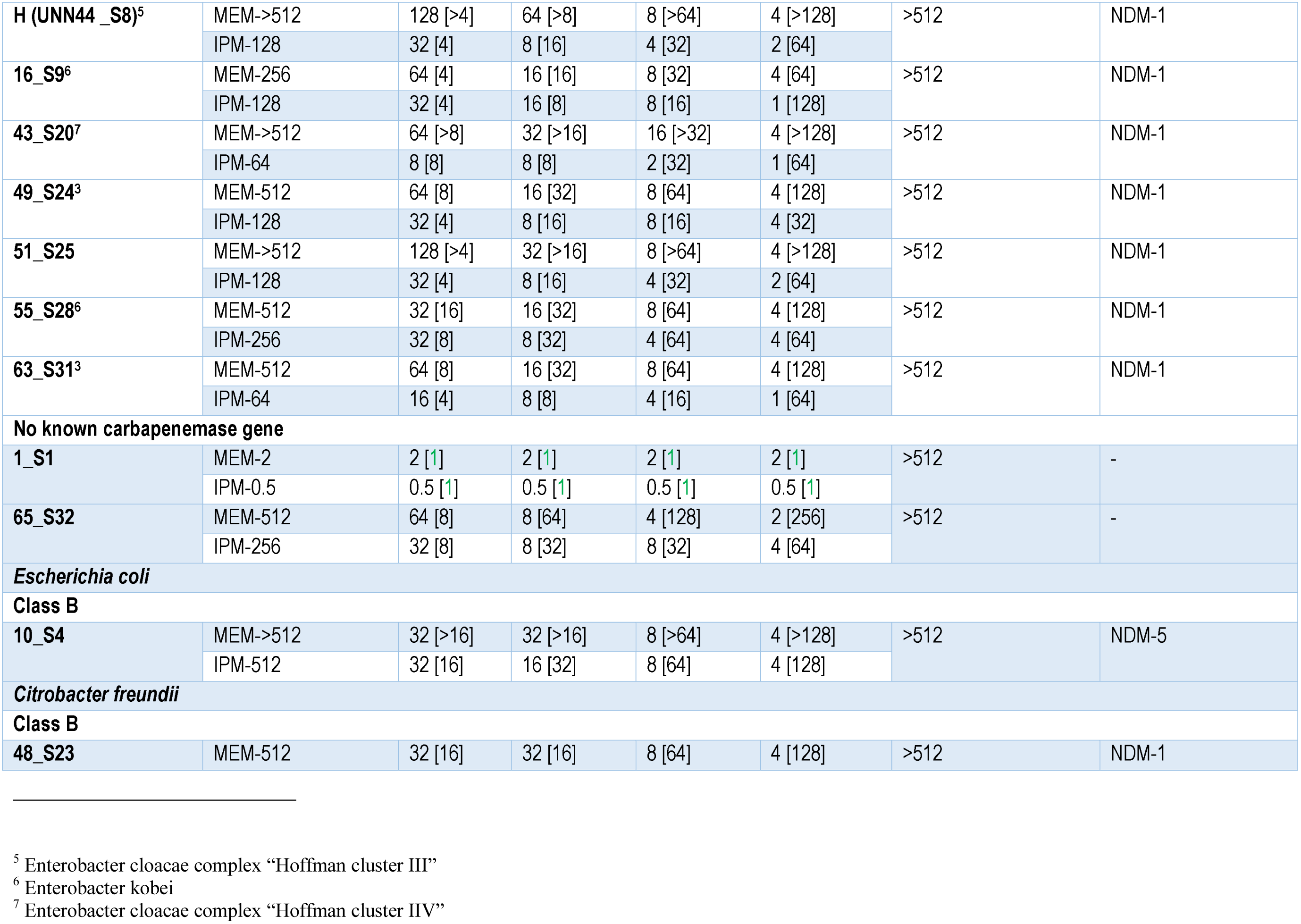
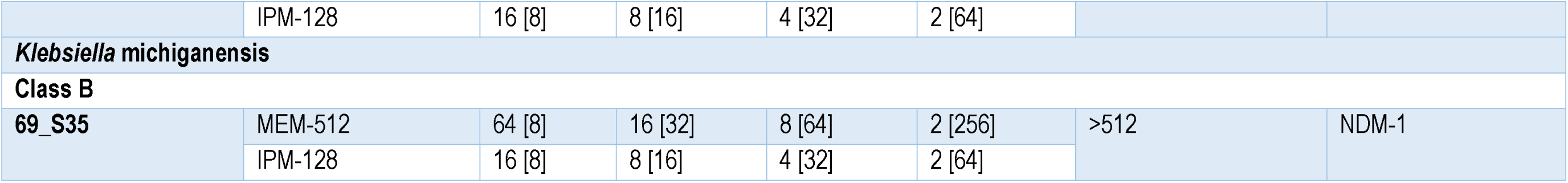
Minimum inhibitory concentrations (MIC) of meropenem (MEM), imipenem (IPM) and Tannic acid (TA) as well as of MEM and IPM in the presence of TA for the clinical South African isolates.

**Table 2:** Minimum inhibitory concentrations (MIC) of meropenem (MEM), imipenem (IPM) and Tannic acid (TA) as well as of MEM and IMP in the presence of TA for the reference isolates.

**in the presence of TA for the reference isolates**
| Isolates | MIC (mg/L) [fold change]<sup>8</sup> |  |  |  |  |  |
| --- | --- | --- | --- | --- | --- | --- |
|  | Antibiotics alone | + Tannic acid |  |  |  | Tannic acid alone |
|  |  | 64 mg/L | 128 mg/L | 256 mg/L | 512 mg/L |  |
| Control |  |  |  |  |  |  |
| *E. coli* ATCC 25922 | MEM- 0.0075 | 0.0075 [1] | 0.0075 [1] | 0.0075 [1] | 0.0075 [1] | >512 |
|  | IMP- 0.06 | 0.06 [1] | 0.06 [1] | 0.06 [1] | 0.06 [1] |  |
| CLASS A |  |  |  |  |  |  |
| KPC-2 |  |  |  |  |  |  |
| *E. coli* KPC-2 | MEM- 16 | 8 [2] | 8 [2] | 8 [2] | 4 [4] | >512 |
|  | IMP- 64 | 8 [8] | 8 [8] | 8 [8] | 8 [8] |  |
| *E. cloacae* KPC-2 | MEM- 32 | 16 [2] | 16 [2] | 16 [2] | 16 [2] | >512 |
|  | IMP- 128 | 64 [2] | 64 [2] | 64 [2] | 32 [4] |  |
| *C. freundii* KPC-2 | MEM- 16 | 4 [4] | 4 [4] | 4 [4] | 4 [4] | >512 |
|  | IMP- 64 | 16 [4] | 16 [4] | 16 [4] | 16 [4] |  |
| *S. marcescens* KPC-2 | MEM- >512 | 512 [≥1] | 512 [≥1] | 512 [≥1] | 512 [≥1] | >512 |
|  | IMP- 512 | 512 [≥1] | 512 [1] | 256 [2] | 256 [2] |  |

<sup>8</sup> A fold change of ≥4 was defined as significant for TA inhibition. Fold changes below 4 are coloured green.

|  |  |  |  |  |  |  |
| --- | --- | --- | --- | --- | --- | --- |
| GES-5 |  |  |  |  |  |  |
| E. cloacae GES-5 | MEM- 64 | 64 [1] | 64 [1] | 64 [1] | 64 [1] | >512 |
|  | IMP- 128 | 128 [1] | 128 [1] | 128 [1] | 128 [1] |  |
| IMI-1 |  |  |  |  |  |  |
| E. asburiae IMI-1 | MEM- 256 | 128 [2] | 32 [8] | 32 [8] | 32 [8] | >512 |
|  | IMP- >512 | 512 [≥1] | 512 [≥1] | 256 [≥2] | 256 [≥2] |  |
| SME-1 |  |  |  |  |  |  |
| S. marcescens SME-1 | MEM- 512 | 256 [2] | 32 [16] | 32 [16] | 32 [16] | >512 |
|  | IMP- >512 | 512 [≥1] | 256 [≥2] | 256 [≥2] | 256 [≥2] |  |
| SME-2 |  |  |  |  |  |  |
| S. marcescens SME-2 | MEM- 512 | 256 [2] | 32 [16] | 32 [16] | 16 [32] | >512 |
|  | IMP->512 | 512 [≥1] | 256 [≥2] | 256 [≥2] | 128 [>4] |  |
| CLASS D |  |  |  |  |  |  |
| OXA-48/181 |  |  |  |  |  |  |
| E. coli OXA-48 | MEM- 1 | 1 [1] | 1 [1] | 1 [1] | 1 [1] | >512 |
|  | IMP- 2 | 2 [1] | 2 [1] | 2 [1] | 2 [1] |  |
| K. pneumoniae OXA-48 | MEM- 8 | 4 [2] | 4 [2] | 4 [2] | 4 [2] | >512 |
|  | IMP- 8 | 4 [2] | 4 [2] | 4 [2] | 4 [2] |  |
| K. pneumoniae OXA-181 | MEM- 1 | 1 [1] | 1 [1] | 1 [1] | 1 [1] | >512 |
|  | IMP- 4 | 4 [1] | 4 [1] | 4 [1] | 4 [1] |  |
| P. rettgeri OXA-181 | MEM- 0.5 | 0.5 [1] | 0.5 [1] | 0.5 [1] | 0.5 [1] | >512 |
|  | IMP- 1 | 1 [1] | 1 [1] | 1 [1] | 1 [1] |  |
| CLASS B |  |  |  |  |  |  |
| NDM-1/NDM-4 |  |  |  |  |  |  |
| E. coli NDM-1 | MEM- 64 | 16 [4] | 8 [8] | 4 [16] | 1 [64] | >512 |
|  | IMP- 128 | 32 [4] | 8 [16] | 1 [128] | 0.5 [512] |  |
| E. cloacae NDM-1 | MEM- 16 | 4 [4] | 4 [4] | 1 [16] | 0.5 [32] | >512 |
|  | IMP- 32 | 8 [4] | 4 [8] | 0.5 [64] | 0.25 [128] |  |
| K. pneumoniae NDM-1 | MEM- 512 | 64 [8] | 16 [32] | 4 [128] | 2 [256] | >512 |
|  | IMP- 256 | 64 [4] | 32 [8] | 16 [16] | 4 [64] |  |
| C. freundii NDM-1 | MEM- 16 | 4 [4] | 2 [8] | 0.5 [32] | 0.5 [32] | >512 |
|  | IMP- 8 | 2 [4] | 2 [4] | 0.5 [16] | 0.25 [32] |  |
| E. coli NDM-4 | MEM- 256 | 32 [8] | 8 [32] | 4 [64] | 2 [128] | >512 |
|  | IMP- 128 | 32 [4] | 16 [8] | 8 [16] | 2 [64] |  |
| VIM-1/19 |  |  |  |  |  |  |
| E. coli VIM-1 | MEM- 64 | 4 [16] | 2 [32] | 0.5 [128] | 0.5 [128] | >512 |
|  | IMP- 128 | 4 [32] | 2 [64] | 1 [128] | 0.5 [512] |  |
| E. cloacae VIM-1 | MEM- 8 | 2 [4] | 2 [4] | 0.5 [16] | 0.5 [16] | >512 |
|  | IMP- 16 | 4 [4] | 2 [8] | 1 [16] | 0.5 [32] |  |
| K. pneumoniae VIM-19 | MEM- 64 | 16 [4] | 8 [8] | 2 [32] | 1 [64] | >512 |
|  | IMP- 512 | 64 [8] | 32 [16] | 4 [128] | 2 [256] |  |
| IMP-1/IMP-8 |  |  |  |  |  |  |
| E. cloacae IMP-1 | MEM- 64 | 16 [4] | 8 [8] | 4 [16] | 1 [64] | >512 |
|  | IMP- 128 | 32 [4] | 16 [8] | 8 [16] | 4 [32] |  |
| E. coli IMP-1 | MEM- 16 | 4 [4] | 2 [8] | 2 [8] | 1 [16] | >512 |
|  | IMP- 32 | 8 [4] | 4 [8] | 4 [8] | 2 [16] |  |
| S. marcescens IMP-1 | MEM- 128 | 64 [2] | 64 [2] | 64 [2] | 32 [2] | >512 |
|  | IMP- 128 | 64 [2] | 64 [2] | 32 [4] | 16 [8] |  |
| E. coli IMP-8 | MEM- 16 | 8 [2] | 4 [4] | 4 [4] | 2 [8] | >512 |
|  | IMP- 32 | 16 [2] | 16 [2] | 16 [2] | 8 [2] |  |
| E. cloacae IMP-8 | MEM- 8 | 4 [2] | 2 [4] | 2 [4] | 1 [8] | >512 |
|  | IMP- 16 | 8 [2] | 8 [2] | 8 [2] | 4 [4] |  |
| K. pneumoniae IMP-8 | MEM- 16 | 8 [2] | 8 [2] | 4 [4] | 1 [16] | >512 |
|  | IMP- 64 | 16 [4] | 16 [4] | 8 [8] | 8 [8] |  |

**Table 3.**
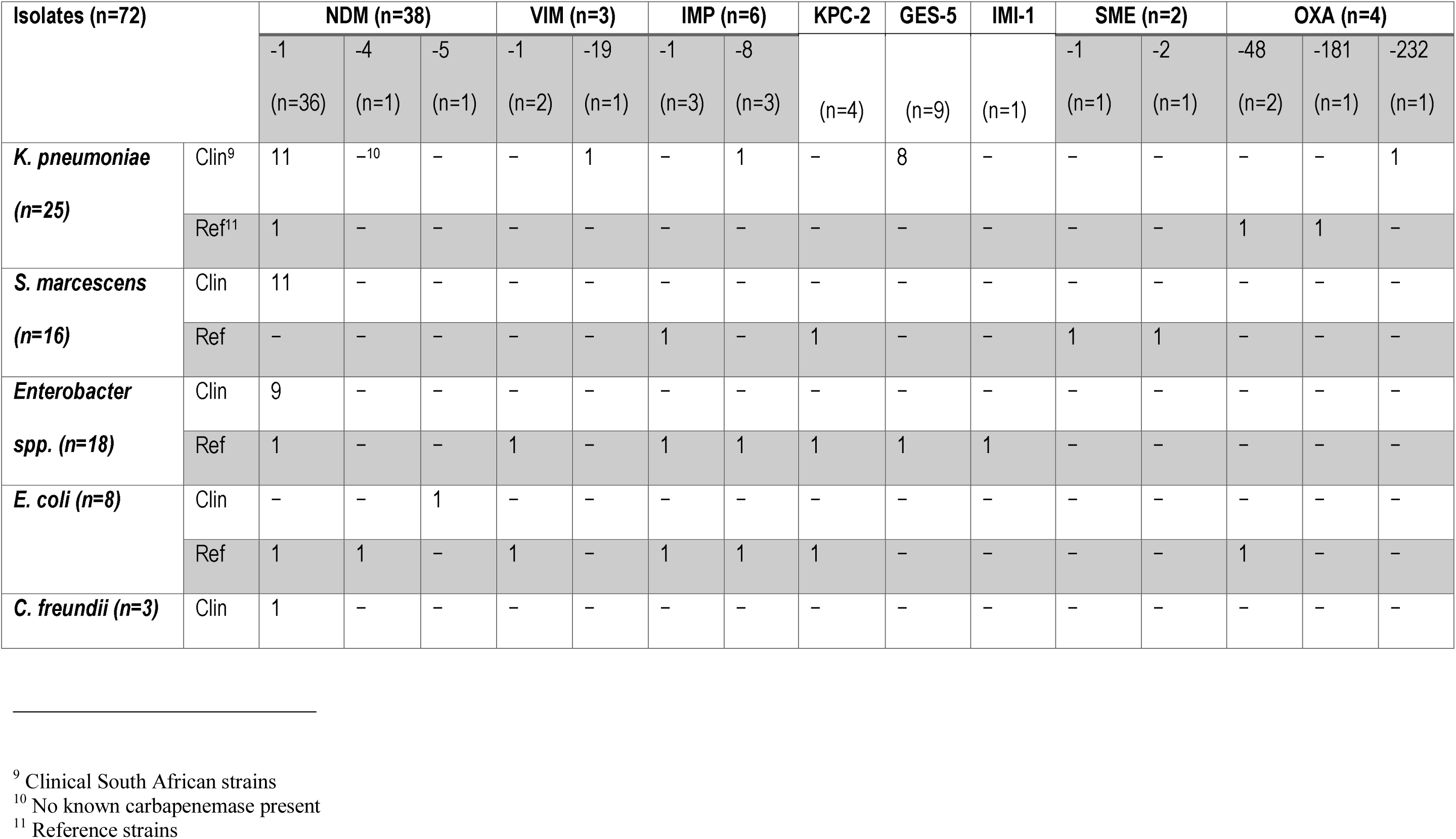
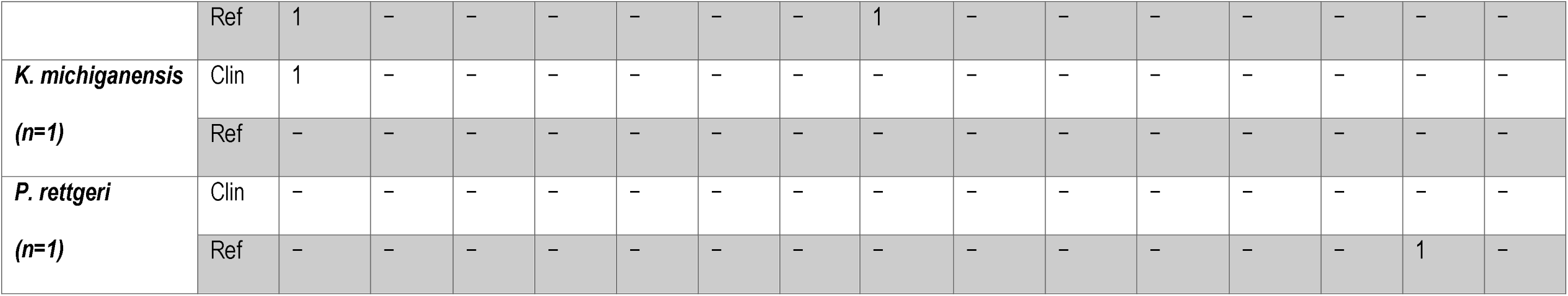
Frequency of carbapenemases found in each Enterobacteriaceae species used in the study.

### 2.2. MICs of CCCP, EPIs and TA. Checkerboard assay of TA

The MICs of RSP, VRP, TZ, CPZ, and CCCP were determined for all the 72 isolates and the control (Tables S1-S4) to determine their innate toxicity to the isolates using protocols already described by the CLSI (Clinical and Laboratory Standards Institute. 2017; Sekyere & Amoako 2017). These MICs were used as a guide in choosing the sub-MICs (i.e. ½ MIC) of these compounds for further testing with MEM and IPM. A sub-MIC was chosen to limit the possibility of the inhibitors killing the cells and interfering with the actual value of the antibiotic-inhibitor combination.

The MIC of TA alone was also determined for all isolates and the control to determine its activity to the isolates according to previously described method and CLSI guideline (Clinical and Laboratory Standards Institute. 2017; Sekyere & Amoako 2017). The checkerboard method was used to evaluate the effect of different concentrations of TA on the carbapenems as described previously (Tascini et al. 2013; King, Sarah A Reid-Yu, et al. 2014). Concentrations of 64, 128, 256, and 512mg/L of TA were added to increasing concentrations of IPM and MEM to determine the MICs of IPM and MEM in the presence of TA (Tables 1-2). The antibacterial assays (MICs) were performed in triplicate.

### 2.3. Role of efflux pumps in carbapenem resistance and in TA’s activity

A sub-MIC of RSP, VRP, TZ, CPZ, and CCCP (Table S3-S4) was used in addition to increasing concentrations of MEM and IMP to evaluate the effects of the former on the MICs of MEM and IMP; a detailed description of this protocol is already described (Osei Sekyere & Amoako 2017; Sekyere & Amoako 2017). This was done to determine whether efflux pumps are involved in TA’s reduction of IMP and MEM MICs.

### 2.4. IC_50_ enzyme inhibition assay

To confirm whether TA has the ability to inhibit the activity of NDM-1 and KPC-2, an *in vitro* enzyme inhibition assay was conducted using purified enzymes that was purchased from Raybiotech (Norcross, GA 30092, USA) and the colorimetric β-lactamase substrate nitrocefin, as previously described (Siemann et al. 2002). Briefly, NDM-1 and KPC-2 enzymes (4nM and 5nM respectively) in HEPES with a pH adjusted to 7.3 ± 0.3 were mixed with 30 μM nitrocefin after 10 to 15 minutes pre-incubation with TA at 30°C. The enzymes were supplemented with 100 μM of ZnCl_2_ and 0.1mg/L bovine serum albumin (BSA). The use of BSA was to minimize the denaturation of the enzymes during experiments. Assays were read in 96 well microtiter plate at 490 nm using a plate reader (SPECTROstar^Nano^ BMG Labtech, Germany) at 25°C. Enzyme assays were conducted in triplicate.

### 2.5. Molecular modelling

The crystal structures (PDB ID: 3QX6, 3RXW, 5ACU and 5FAS) were retrieved from the RSCB Protein Data Bank (https://www.rcsb.org/pdb/). The missing residues were added using a graphical user interface of Chimera, a molecular modelling tool (Pettersen et al. 2004). A ligand interaction map was generated using the web version of PoseView (Stierand & Rarey 2010). System preparation, molecular docking and molecular dynamic simulations where carried out to ascertain the interactions of the enzymes (NDM-1, VIM-2, KPC-2) with the ligand (TA). Details of the procedure used has been stated in the supplementary document.

### 2.6. Cytotoxicity Assay

HepG2 cells were maintained at 37°C in 10% complete culture medium (CCM; Eagles minimum essential medium supplemented with foetal bovine serum, antibiotics, and L-glutamine) until confluent. Following trypsinization, 15000 cells/well (300μl) were allowed to adhere to a 96-well plate overnight. The cells were treated with seven dilutions of TA (16 - 1024mg/L) prepared in CCM; untreated cells (CCM only) served as the control. After 24 hours the treatment was replaced with 20μl MTT solution (5mg/mL MTT in PBS) and 100μl CCM for 4 hours. The resulting formazan product was solubilized in 100μl DMSO (1 hour) and the absorbance at 570nm/690nm was determined (BioTek μQuant plate reader; BioTek Instruments Inc., USA). The average absorbance values performed in triplicate were used to calculate cell viability.

### 2.7. Phylogenetic analysis

Fifteen carbapenemase genes were found in all the 72 isolates included in this study. Reference amino acid sequences of these 15 carbapenemase genes viz., NDM-1, NDM-4, NDM-5, IMP-1, IMP-8, VIM-1, VIM-19, KPC-2, GES-5, IMI-1, SME-1, SME-2, OXA-48, OXA-181, and OXA-232 were downloaded from GenBank and aligned with MAFFT online server (http://mafft.cbrc.ip/alignment/server/). The phylogenetic relationship between the carbapenemases was determined using the MAFFT phylogeny server (http://mafft.cbrc.ip/alignment/server/phvlogeny.html). The phylogenetic tree was viewed online and downloaded with Newick internal labels (Fig. 2). This was carried out to offer possible insights into the potentiating differences of TA on the different carbapenemase enzymes.

### 2.8. Data and statistical analysis

The frequency of the carbapenemases per Enterobacteriaceae species was determined from Tables 1 and 2 by calculating the number of isolates expressing a particular enzyme (Table 3). The MIC fold change (Δ), defined as the ratio of the MIC of MEM or IMP alone to that of TA plus MEM or IMP, was calculated for each isolate and shown in a square bracket in Tables 1 and 2. A fold change (Δ) of ≥4 was adopted as significant and all Δ below 4 are coloured green in Tables 1 and 2. A mean fold change per antibiotic was calculated for every enzyme (Table 3) using the following equation:

Sum (Δ of all isolates expressing x)/Total number of isolates expressing x

Where *x* is any of the carbapenemases found in this study’s isolates. The mean fold changes per antibiotic and carbapenemase was translated into a bar graph using Microsoft Excel to show the relative effect of TA (512 mg/L) on each carbapenemase (Fig. 1).

**Fig. 1:**
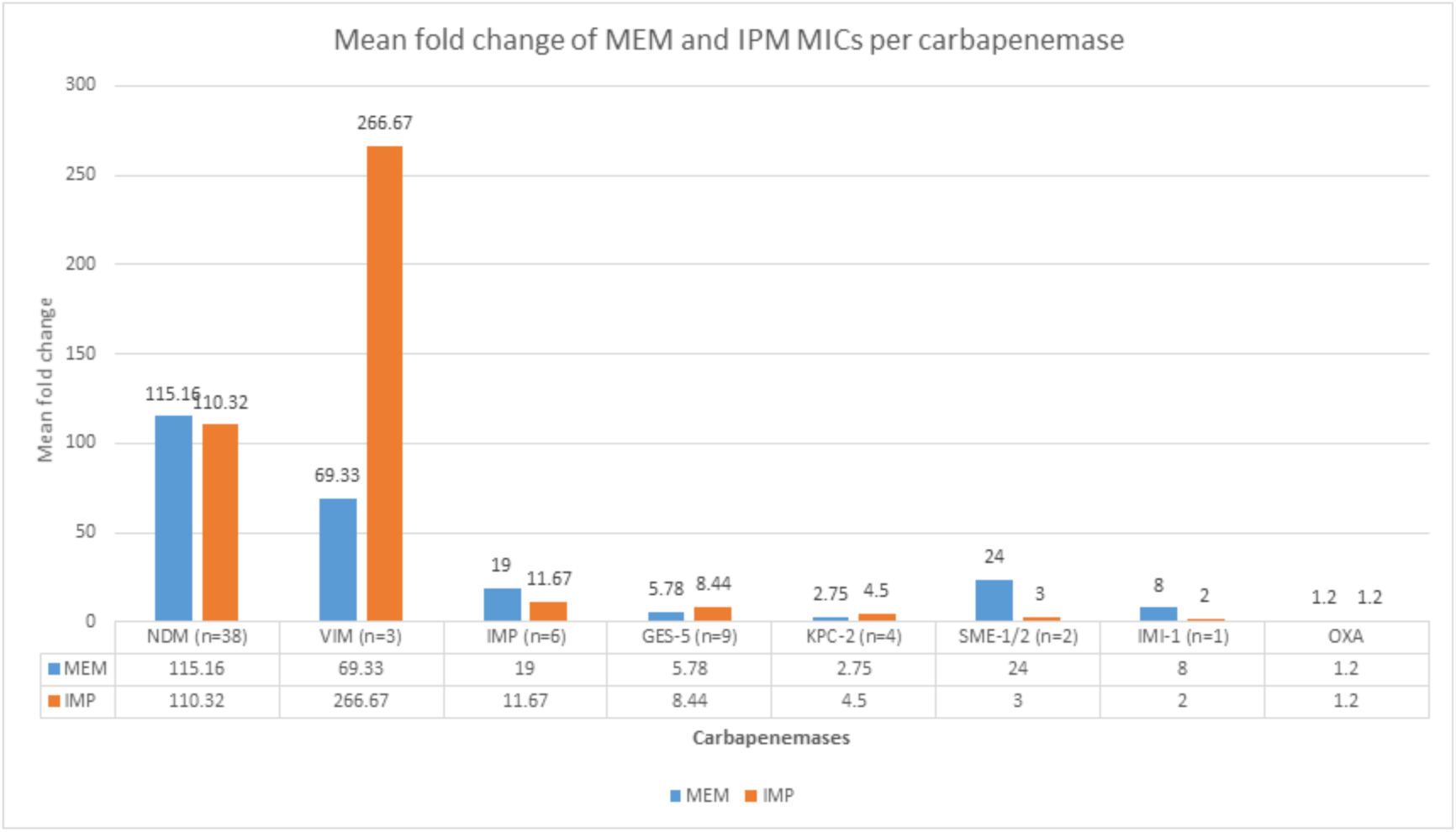
MEM and IPM MIC fold change of isolates per carbapenemase upon addition of tannic acid (TA). The effect of tannic acid (512mg/L) in reducing and/or reversing meropenem (MEM) and imipenem (IPM) resistance among Enterobacteriaceae is shown above as a mean fold change (Δ). The mean Δ is measure of the relative effect of TA on the MIC of each isolate. TA effect was most potent against bacteria producing metallo-β-lactamases such as NDM (MEM=115, IPM=110), VIM (MEM=69, IPM=266) and IMP (MEM=19, IPM=12) than against those producing serine-β-lactamases such as GES (MEM=6, IPM=8), KPC (MEM=3, IPM=5), SME (MEM=24, IPM=3) and IMI (MEM=8, IPM=2); these effects were statistically significant (p-value < 0.0001-0.01). Effect on producers of OXA-type carbapenemases (MEM=1, IPM=1) was very minimal or null and statistically insignificant (p-value > 0.01). IMP and MRP was thus affected differently by TA per enzyme.

GraphPad Prism 5.0 (GraphPad Software, San Diego, CA, USA) was used to determine the IC_50_, the significance of the Δ and mean Δ per antibiotics per isolate/species and per expressed carbapenemase in the presence of the efflux pump inhibitors and TA (Tables S5-S6). MIC fold changes (Δ) with a P-value of <0.01 were considered statistically significant.

## 3. Results

### 3.1. TA significantly reduces imipenem (IPM) and meropenem (MEM) MICs in classes A and B carbapeneamase-producing isolates

The checkerboard method was used for this assay. Increasing concentrations of TA resulted in decreasing imipenem (IPM) and meropenem (MEM) MICs for particularly NDM-, VIM-, IMP-, GES-5-, SME-1/-2-, IMI-1-, and KPC-2-producing isolates in a descending order of magnitude (Δ) (Tables 1-2; Fig. 1). TA reversed resistance to IPM and MEM in all MBL-positive isolates (p-value < 0.0001-0.01) except in one *Serratia marcescens bla*_IMP-1_- positive strain. On the contrary, TA significantly (p-value < 0.001-0.01) reduced the MEM MICs of most isolates expressing IMI and SME carbapenemases whilst it reversed resistance to only a few KPC-2- and GES-5-positive isolates. The effect of TA on OXA-48/-181/-232-producing isolates was insignificant (p-value > 0.01) although it halved the MIC of one *Klebsiella pneumoniae* OXA-48-positive strain (Tables 1-2; Fig. 1). TA’s resistance-modulating effect was most potent against producers of MBLs such as NDM, VIM, and IMP than against producers of serine-β-lactamases such as GES, KPC, SME, and IMI. The effect of TA on the tested isolates was greatest at a concentration of 512mg/L for all the carbapenemases and was thus used in calculating the mean Δ (Fig. 1); unless otherwise stated, the effects of TA on the CRE in this manuscript are discussed with the premise that a concentration of 512mg/L was used. TA on its own had no effect or activity on the isolates even at >512mg/L, whilst CCCP and the other EPIs exhibited inhibition to all the tested organisms (Tables 1-2, and Tables S1-S2).

### 3.2. Efflux pump inhibitors and CCCP fail to reduce MEM and IMP MICs significantly

In order to investigate whether TA reduces MICs of MEM and IMP through efflux pumps inhibition, EPIs and CCCP were employed to elucidate this hypothesis. As shown in Tables S1-S2, all the EPIs and CCCP, which is known to indirectly block efflux activity through reduction in both adenosine triphosphate (ATP) production and the proton motive force (PMF),(Sekyere & Amoako 2017) failed to significantly reduce or reverse resistance to MEM and IMP in all but one *Serratia marcescens* isolate, B (UNN38 _S2) (Table S1). This indicated that TA did not reduce the MICs of MEM and IPM through inhibition of efflux pumps activity as reported in methicillin-resistant *Staphylococcus aureus* (MRSA) strains (Myint et al. 2013).

### 3.3. Enzymatic assay and molecular modelelling and docking

Inhibition of efflux pumps not being involved in the protection of MEM and IMP, we therefore investigated the interaction between carbapenemase enzymes and TA to evaluate the potentiating effect of TA toward MEM and IMP. Half-maximum inhibitory concentration (IC_50_) values of TA were determined using nitrocefin for NDM-1 (MBL) and KPC-2 (SBL) to determine the difference in inhibition between the two classes. TA respectively inhibited ≈50% of NDM-1 and KPC-2 at 15 μM and 270 μM indicating that TA possess an effect on both classes of carbapenemase enzymes.

Furthermore, computational molecular dynamic analyses were employed to ascertain that TA interacts with class A and class B carbapenemase enzymes as proof of concept. Thus, root-mean-square deviation (RSMD) analysis verified that all four tested systems (ligand-enzymes complexes: TA-_NDM-1_, TA-_VIM-2,_ TA-_KPC-2_ TA-_OXA-48_) were quite stable over the molecular docking simulations and that, the interactions with the ligand site were strong enough to maintain the latter bonding (Fig. 2A, and Table S7). The radius of gyration (Rg) of the four systems were also steady, highlighting the moderate conformational changes occurring during the simulation (Fig. 2B). These results demonstrated that TA interact with each enzyme by reducing their hydrolysis activities at different levels, and also showed a stable compact structure. However, OXA-48 and KPC-2 (serine-β-lactamases) exhibited a significant overall high radius of gyration in comparison to the metallo-β-lactamases (VIM-2 and NDM-1) (Fig. 2B). The difference(s) in the interactions between the catalytic active site residue(s) and TA amplified the conformational flexibility of the complex and ultimately affected the receptor grip on the inhibitor, resulting in a higher radius of gyration of the SBLs. Further analysis of the binding free energy showed that the MBLs (VIM-2 and NDM-1) exhibited a higher binding affinity when compared to the SBLs. These findings corroborate the results from the Rg and the *in vitro* experiments. Analysis of ligand-enzymes interaction (Fig. 3) showed that the MBLs interact with a larger number of their active site residues as opposed to that of OXA-48 and KPC-2. TA also forms electrostatic interactions with zinc.

**Fig. 2:**
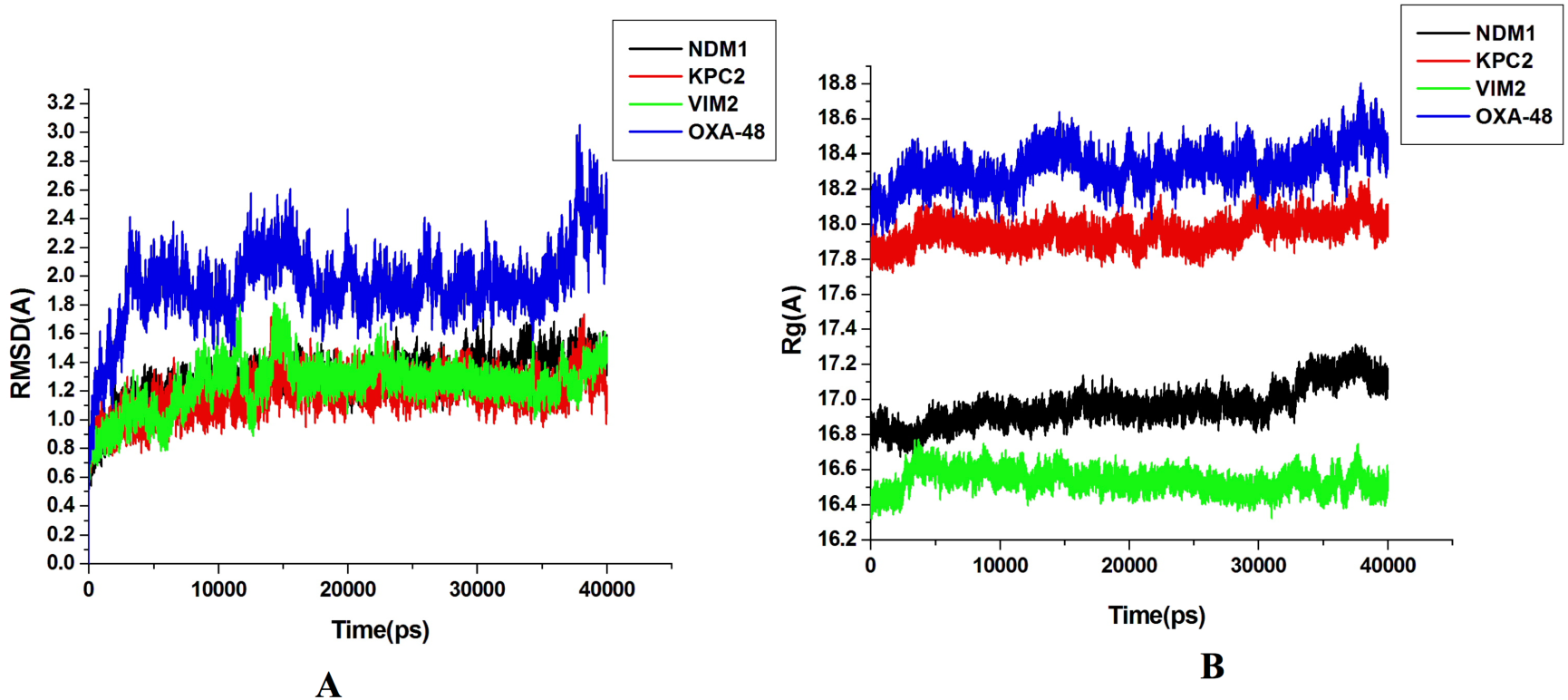
The Root-mean-square deviation (RMSD) and Radius of gyration (Rg) diagrams of the four complexes: A) RMSD showing the stability of the systems (enzyme-ligand complex); B) Rg illustrating the compactness of the systems.

**Fig. 3:**
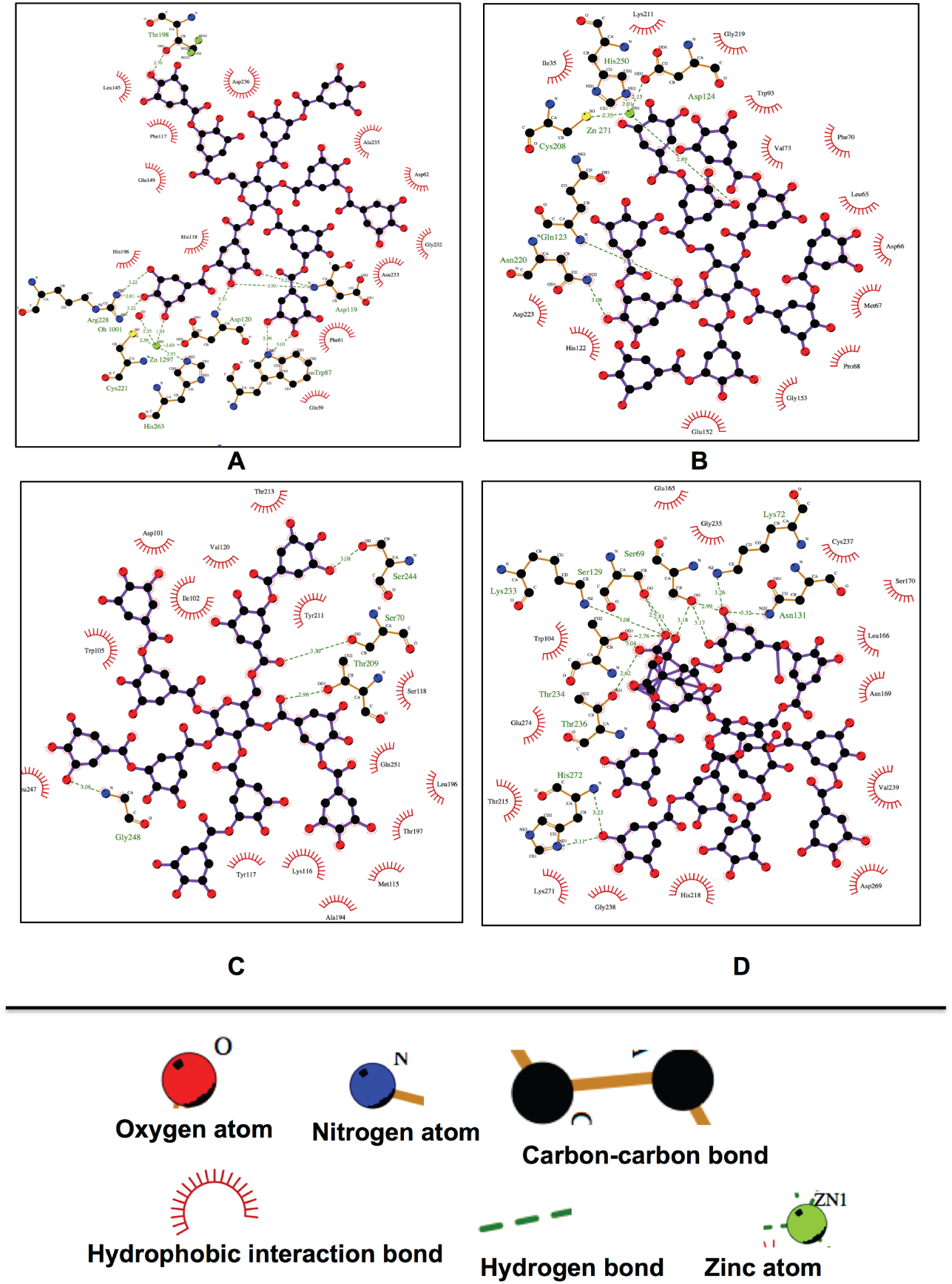
A ligplot showing the relative position of Tannic acid inside the active site of: (A) VIM-2, (B) NDM-1 (C) OXA-48, D) KPC-2. TA interact more with MBLs when compared to SBLs at their catalytic active sites.

### 3.4. Cytotoxicity assay

The MTT assay was used to measure TA cytotoxicity (0 - 1024mg/L) in HepG2 cells after 24 hours exposure. An increase in cell viability was associated with TA concentrations; 16mg/L (118.98±6.95%) and 32mg/L (108.52±7.28%) (Fig S2). However, for the 64mg/L, 128 mg/L and 256 mg/L the cell viability decreased to 89.26±8.37%, 63.08±4.82% and 63,40±4.73% respectively. The metabolic activity of TA was increased again at 512mg/L (93.50±4.86%) and 1024mg/L (93.86±4.43%) (Fig S2). TA was having tolerable cytotoxic effect for all the concentrations tested.

### 3.5. Phylogenetic tree of the carbapenemases

A phylogenetic tree of all the carbapenemases (n=15) was drawn to ascertain offer plausible insights into the potentiating effects on the various carbapenemases. The tree (Fig. 4) shows that all the members of the same carbapenemase class are clustered closer together (or share closer sequence identity) and separated from members of other classes that were found on different branches. MBLs (NDM, VIM and IMP) were clustered together and farther from the other subclasses. GES-5 was more distant from KPC-2, IMI-1 and SME whilst IMI-1 and SME were very closely related and distant from KPC-2. All the three OXA-type carbapenemases had a closer sequence similarity as shown by their close clustering on the same branch (Fig. 1 and 4).

**Fig. 4:**
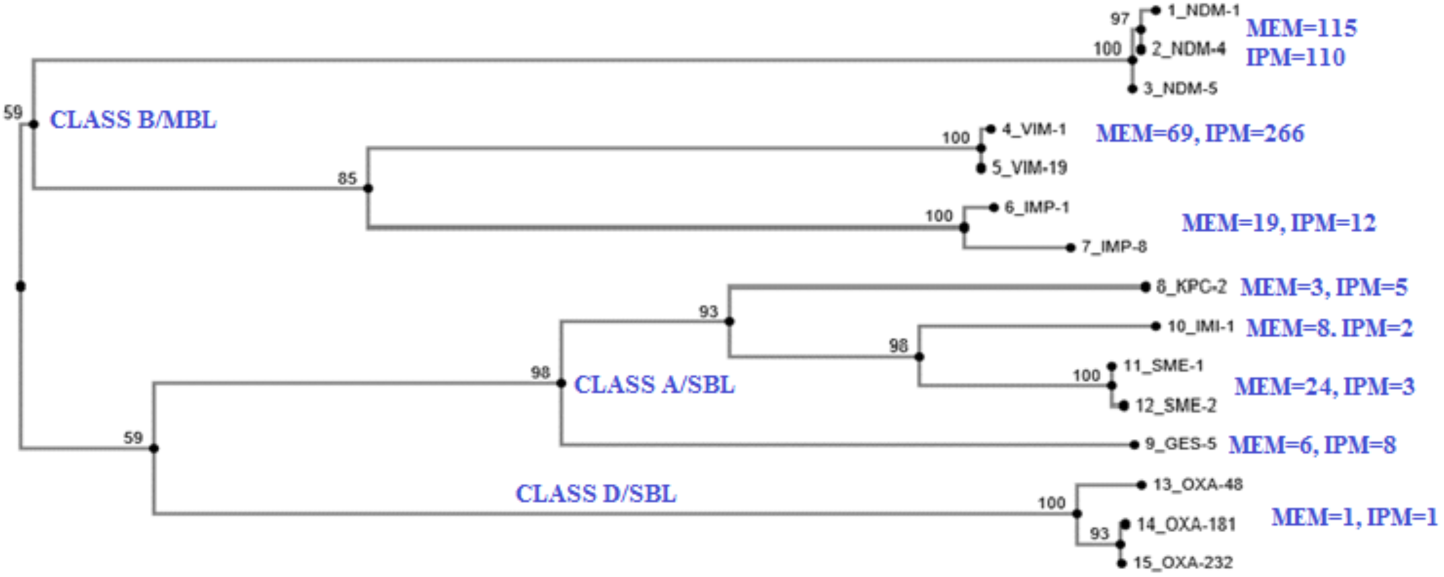
Phylogenetic tree of the carbapenemases. The carbapenemases found in the isolates used in this study are clustered into their families of classes A, B, and D, showing the close relationship and amino acid sequence identity between members of the same family. The mean MIC fold change per enzyme and antibiotic (MEM and IMP) are shown besides the enzyme names. The effect of TA differed from carbapenemase to carbapenemase and from carbapenemase class to class, with class B being the most inhibited.

### 3.6. Species, carbapenemases and mean fold change

The study comprised of Enterobacteriaceae isolates from six genera and 12 species expressing eight carbapenemases, namely NDM (n=38), IMP (n=6), VIM (n=3), GES-5 (n=9), KPC-2 (n=4), SME (n=2), IMI-1 (n=1), and OXA-48-types (n=5) (Tables 1-2). Three clinical isolates from South Africa i.e. 1-S1, 65_S32, and 45_S21 were non-CPEs and had tested negative both for the Carba-NP test (Osei Sekyere et al. 2015; Osei Sekyere, Govinden, & Essack 2016) and efflux-mediated carbapenem resistance (as shown in the MIC fold change of 1), but the latter two isolates had very high IPM and MEM MICs, which were drastically reduced by TA (Table 1). Of note, isolate 3_S2 was the only one that expressed OXA-232 with an IPM and MEM MIC of 128mg/L; however, it was Carba-NP negative (Osei Sekyere, Govinden, & Essack 2016).

The mean Δ of IPM and MEM for each carbapenemase was calculated from the individual Δ of all the isolates to serve as an index of the relative effect of TA on each carbapenemase (Fig. 1). Thus, carbapenemases with higher mean Δ were most affected by TA. Disparities were also observed between the mean Δ of IPM and MEM for all the carbapenemases. The highest mean Δ for IPM and MEM were observed in isolates expressing NDM, VIM and IMP followed by those expressing GES-5, SME, KPC-2, and IMI-1. OXA-48-type-expressing isolates had the least mean Δ for the two antibiotics (Fig. 1 and 4).

## 4. Discussion

Carbapenems have become the last-resort antibiotic for treating fatal and multidrug-resistant bacterial infections due to increased fluoroquinolone, aminoglycoside and cephalosporin resistance (Osei Sekyere, Govinden, & Essack 2016; Osei Sekyere 2016; Osei Sekyere & Amoako 2017). The increased use of carbapenems to battle these resistant infections have thus led to the selection and dissemination of CPE, which are resistant to almost all known β-lactam antibiotics (Nordmann & Poirel 2014; Sekyere et al. 2016). In this study, we report on a plant-derived agent, TA, which protect carbapenem antibiotics from enzymatic hydrolysis by inhibiting the activity of mostly class B MBLs and class A SBLs, thus, reducing carbapenem MICs and resistance.

The inability of CCCP and the other EPIs to reverse IPM and MEM resistance suggests that TA does not inhibit efflux pumps to reverse IPM and MEM MICs. Moreover, TA alone had no biocidal effect on the isolates at concentrations >512 mg/L, as seen from the MIC of TA (Tables 1-2), ruling out the possibility that TA’s mechanism of action is through protein inhibition. As well, TA does not interact negatively with the carbapenems, as such an interaction would not have produced the results seen. Rather, a combination of TA with IPM and MEM resulted in a reduction or modulation of carbapenem resistance in most isolates, thus revealing the potentiating property of TA toward both tested carbapenem antibiotics.

Notably, the effect of TA was mainly dependent on the type of carbapenemase enzyme expressed by the isolates rather than by the species or strain type. For instance, irrespective of the species or strain, TA modulated carbapenem resistance in all MBL-producing isolates as well as in most isolates expressing certain types of class A SBLs. However, resistance in the same species expressing the same class D carbapenemase types were barely modulated. Hence, the resistance-modulating effect of TA was mainly based on the type of carbapenemase rather than on the species or strain, indicating that TA interacts with the carbapenemase enzymes to modulate carbapenem resistance, similarly to previous studies that demonstrated inhibition of carbapenemase enzymes to improve carbapenem antibiotics such as aspergillomarasmine A, 1,4,7-triazacyclononane- 1,4,7-triacetic acid (NOTA), di-(2-picolyl)amine (DPA) but with broader spectrum of activity (King, Sarah A. Reid-Yu, et al. 2014; Somboro et al. 2014; Azumah et al. 2016). To confirm this hypothesis, *in vitro* enzyme inhibition assays and computational molecular analysis were conducted. The enzyme assay showed a wide difference in TA inhibition capacity per carbapenemase. The IC_50_ values further affirmed that TA inhibits MBLs more prominently than SBLs. Computational studies predicted that TA inhibit both MBLs and SBLs by targeting their hydrophobic sites. Analysis of ligand- enzymes interaction indicated that the MBLs interact with a larger number of their active site residues as opposed to that of OXA-48 and KPC-2 (SBL). TA also forms hydrogen bonds with zinc, which is important for MBL activity. This binding decreases the enzyme activity harming or killing the bacteria. Further analysis of the binding free energy showed that the MBLs (VIM-2 and NDM-1) exhibited a stronger and stable binding affinity when compared to the SBLs, which is caused by the differences in the interactions between the catalytic active site residue(s) and TA, amplifying the conformational flexibility of the complex and ultimately affecting the receptor grip on the inhibitor (Table 4). This results in a higher radius of gyration of the SBLs than MBLs. These findings corroborate the results from the *in vitro* experiments and confirmed that TA interacts with the enzymes to potentiate IMP and MEM efficacy. The per-residue interaction energy decomposition calculations applied in this study will potentially assist medicinal chemists in the design of inhibitors that can interact with these carbapenemase residues in future studies.

**Table 4:** MM/GBSA binding free energy profile of all four systems. All energy terms are presented in kcal/mol.

| Systems* | Evdw | Eelec | $\Delta G_{solv}$ | $\Delta G_{gas}$ | $\Delta G_{bind}$ |
| --- | --- | --- | --- | --- | --- |
| **MBLs** |  |  |  |  |  |
| NDM-1 | -3.889±0.1660 | -34.6186±0.3174 | -38.5080±0.2285 | -5.7249±0.1561 | -44.2329±0.3806 |
| VIM-2 | -3.90744±0.1014 | -31.4537±0.1706 | -35.3612±0.1070 | -8.3608±0.1485 | -43.7220±0.4513 |
| **SBLs** |  |  |  |  |  |
| KPC-2 | -1.9446 ±0.1870 | -17.3093±0.2722 | -19.2540±0.1590 | -2.8624±0.0752 | -22.1164±0.0111 |
| OXA-48 | -1.0841±0.1321 | -17.9093±0.1887 | -18.9935±0.1818 | -3.1591±0.1712 | -22.5275±0.1300 |
| $\Delta E_{vdw}$, van der Waals energy; $\Delta E_{elec}$, electrostatic energy; $\Delta G_{solv}$, polar solvation energy; $\Delta G_{gas}$, nonpolar solvation energy; $\Delta G_{bind}$, total binding free energy; *Enzyme-ligand complex. | | | | | |

Among carbapenemases, class B is known to have little amino acid homology and longer evolutionary distance to classes A and D as seen on the phylogenetic tree, which shows that class A and D have a closer evolutionary distance (Fig. 2). In addition, NDM is most closely related to VIM than to IMP with a sequence identity of 32.4% (Saini 2012). Hence, the closer amino acid sequence homology of VIM to NDM rather than its closer evolutionary distance to IMP might in part explain the observation in which VIM-positive isolates had greater mean MIC fold changes than IMP- positiveones (Fig. 2)

Isolates 65_S32 and 45_S21 both tested negative for the Carba-NP test and no known carbapenemase was found in their genome although they both had very high IPM and MEM MICs (45_S21=32mg/L; 65_S32=256 and 512mg/L respectively) (Osei Sekyere, Govinden, & Essack 2016; Osei Sekyere & Amoako 2017; Sekyere & Amoako 2017). Furthermore, both isolates had no MIC change when tested with carbapenems-EPIs, indicating that their efflux pumps were not possibly responsible for the high MICs recorded. However, TA could reduce/modulate resistance to carbapenems in these isolates. As the OXA-232-producing isolate, 3_S2, was also Carba-NP test negative and could not be affected by TA, but was highly resistant to IPM and MEM, we suspect that an unknown or undiscovered MBL or class A SBL might be mediating the high carbapenem MIC recorded in these two strains (Table 1).

Available evidence from published studies show that tannins, which includes TA, catechin, ethylgallate, epicatechin gallate, epigallocatechin gallate, methyl gallate, ellagic acid, rosmaric acid and 1,2,3,4,6-Penta-O-galloyl-b-D-glucopyranose are polyphenolics that are abundant in plants (Leal et al. 2011; Slobodníková et al. 2016) (Fig. S1). Tannin extracts and fractions have been reported to either potentiate or modify the effects of β-lactams, fluoroquinolones, and several other antibiotic classes (Leal et al. 2011; Myint et al. 2013; Slobodníková et al. 2016). Moreover, they have been shown to inhibit biofilm formation, quench quorum sensing, and confer antimicrobial and antioxidant activity against bacteria by binding to bacterial proteins, cell wall and cell membrane structures, resulting in cell lysis and efflux pumps inhibition (Leal et al. 2011; Myint et al. 2013; Slobodníková et al. 2016). However, the reported antibacterial and efflux inhibition effects of tannins were not seen in this study, intimating that other tannins besides TA may be responsible for those effects.

In an earlier study, Leal et al. (Leal et al. 2011) found that the tannin fractions of *Pentaclethra macroloba* was bactericidal to bacteria in a dose-dependent manner by inhibiting protein synthesis, although protein binding is the universally accepted mechanism of tannins’ biological and astringent activity (Slobodníková et al. 2016). Notably, the tannin fraction was found to be non-toxic to eukaryotic cells at the effective bactericidal concentration (Leal et al. 2011). As TA alone was not active against the Enterobacteriaceae isolates used in this study it is obvious that it did not inhibit protein synthesis supporting our hypothesis a potential carbapenemase inhibitor.

## 5. Conclusion

TA did not reverse carbapenem resistance by inhibiting all proteins/enzymes and/or efflux pumps in the cells, but through an inhibition of the carbapenemases by a possible binding/sequestration interaction with their catalytic-active sites, thus freeing the carbapenems to exert their antibiotic effect. Further investigation of TA properties might extend its inhibitory spectrum to cover all carbapenemases and reduce the efficacy of carbapenems in carbapenem-resistant Enterobacteriaceae.

## 7. Transparency Declaration

Professor Sabiha Essack is a member of the Global Respiratory Infection Partnership sponsored by an unconditional education grant from Reckitt and Benckiser.

## 8. Author contributions

Co-conceptualized the study: AMS, DGA, JOS, LAB and SYE. Performed the experiments: AMS, JOS, DGA, HMK and RK. Analyzed the data: AMS, JOS, DGA, HMK and RK. Validation of the results: All. Wrote the paper: JOS. Undertook critical revision of the manuscript: All. Supervision: LAB and SYE.

